# T4SEpp: a pipeline integrated with protein language models effectively predicting bacterial type IV secreted effectors

**DOI:** 10.1101/2023.07.01.547179

**Authors:** Yueming Hu, Yejun Wang, Xiaotian Hu, Haoyu Chao, Sida Li, Qinyang Ni, Yanyan Zhu, Yixue Hu, Ziyi Zhao, Ming Chen

## Abstract

Many pathogenic bacteria use type IV secretion systems(T4SSs) to deliver effectors (T4SEs) into the cytoplasm of eukaryotic cells, causeing diseases. The identification of effectors is a crucial step in understanding the mechanisms of bacterial pathogenicity, but this remains a major challenge. In this study, we used the full-length embedding features generated by six pre-trained protein language models to train classifiers predicting T4SEs, and compared their performance. An integrated model T4SEpp was assembled by a module searching full-length, signal sequence and effector domain homologs of known T4SEs, a machine learning module based on the hand-crafted features extracted from the signal sequences, and the third module containing three best-performing protein language pre-trained models. T4SEpp outperformed the other state-of-the-art (SOTA) software tools, achieving ∼0.95 sensitivity at a high specificity of ∼0.99, based on the assessment of an independent testing dataset. Additionally, we performed a comprehensive search among 8,761 bacterial species, leading to the discovery of 227 species belonging to 3 phyla and 117 genera that possess T4SSs. Furthermore, leveraging the power of T4SEpp, we successfully identified a grand total of 12,622 plausible T4SEs. Overall, T4SEpp provides a better solution to assist in the identification of bacterial T4SEs, and facilitates studies of bacterial pathogenicity. T4SEpp is freely accessible at https://bis.zju.edu.cn/T4SEpp.

## Introduction

Gram-negative bacteria employ more than one dozen of secretion systems to transport proteins out of the cell envelope[1, 2]. Among them, the type IV secretion system (T4SS) is a complex molecular machine spanning both the inner and outer membranes, and translocate substrate proteins into eukaryotic host cells in only one step[3-9]. Protein-translocating T4SSs can be divided into two major families according to the composition of component elements: type IVA, exemplified by the *A. tumfaciens* VirB/VirD4 T4SS and *H. pylori* Cag T4SS, and type IVB exemplified by *Legionella* Dot/Icm T4SS[9]. Substrate proteins translocated by T4SSs, also called effectors, play important roles in bacterial infections and pathogenicity[1, 10, 11].

Effectors of T4SSs (T4SEs) are transported directly or as complexes with DNA in many pathogenic bacteria, such as *Helicobacter pylori*, *Legionella pneumophila*, *Bordetella pertussis*, *Coxiella*, *Brucella*, and *Bartonella*[12-17]. T4SS-mediated entry of effector proteins into recipient cells is contact-dependent[18]. Once they enter the eukaryotic host cytoplasm, they disrupt signal transduction and cause various host diseases. Identifying these effectors is crucial for understanding the mechanisms of infection and pathogenicity caused by these bacteria. However, because the composition and sequences vary significantly, it is challenging to identify new T4SEs experimentally. Although many T4SEs have been identified and characterized in a few model organisms[19-22], the exact mechanism remains unclear.

Since 2009 when the first machine-learning algorithm was introduced, tens of computational models have been developed to predict T4SEs[2, 23]. Early algorithms were mainly species-specific, such as those predicting T4SEs in *Legionella pneumophila*[23]. In another study, Wang *et al.* developed an SVM-based model, T4SEpre, which exhibited good overall and cross-species performance[24]. However, T4SEpre only considers the features buried in the C-terminal 100 amino acids[24]. More studies, especially ensemble models recently developed with multi-aspect features, learn features from full-length proteins to improve performance[25, 26]. Deep learning algorithms have also been applied in for the prediction of T4SEs. For example, CNN-T4SE integrated three convolutional neural network (CNN) models to learn the features of amino acid composition, solvent accessibility, and secondary structure of the full-length T4SEs[27]. T4SEfinder is a multi-layer perception (MLP) model that learns the features generated by a pre-trained BERT model[28], which can predict T4SEs accurately[29]. Notably, BERT is a natural language processing (NLP) model that is appealing in biology and other fields[30-35]. NLP models have been successfully applied to the prediction of protein subcellular localization[31, 32], secondary structure[32, 33, 35], and others[34]. Besides T4SEfinder, the NLP-based pre-trained transformers have also been used for the prediction of bacterial type III secreted effectors and Sec/Tat substrates, both achieving superior performance[36, 37].

Although machine learning strategies have achieved some success in the identification of T4SEs[2, 23, 24], the high false-positive rate has been a big challenge. To reduce the false-positive rate in predicting type III effectors, Hui et al. proposed a strategy to combine machine learning models with homology searching, and integrate multiple modules considering the multi-aspect biological features of the effector genes[38]. To improve model performance, other models have also considered the multiple features and a combination of homology-based strategies in the prediction of type III effectors[39-41]. For T4SE prediction, homology searching was also been applied independently. For example, S4TE integrates 13 sequence homology-based features, including homology to known effectors, homology to eukaryotic domains, presence of subcellular localization signals, and secretion signals, and develops a scoring scheme to predict T4SEs mainly from α- and γ-proteobacteria[42]. Despite the high precision, the sensitivity could be influenced by the large diversity of T4SE composition and sequences. Therefore, it could be a better solution to take the advantages of both machine learning approaches, especially ensemblers, and homology-based methods, designing an integrated T4SE prediction pipeline that combines various models and comprehensively considers various characteristics of effector sequences.

In this study, we proposed a hybrid strategy for predicting T4SEs. First, a homology searching strategy scanned both the global homology of full-length proteins and the local homology of domains to known effectors. Additionally, we retrained a machine learning module T4SEpre[24] with updated T4SE data and hand-crafted amino acid composition features in the C-termini. Furthermore, a group of transfer learning models was developed based on the features generated by various pretrained transformers. For the transfer learning models, we utilized the deep context protein language models ESM-1b, ProtBert, ProtT5-XL, and ProtAlbert to represent protein sequence features[32, 33]. These features can characterize the intrinsic but unclear properties of protein sequences and the interactions between positions. Based on these feature representations, application models were developed to classify T4SEs using a deep neural network architecture with an attention mechanism. Finally, we integrated the homology-based modules, machine learning models based on traditional handcrafted features, and transfer learning models with transformer-generated features into a pipeline, namely T4SEpp, which assembles the individual modules in a linear function to generate a prediction score reflecting the likelihood of a protein to be a T4SE. A web application for T4SEpp is also available via the link: https://bis.zju.edu.cn/T4SEpp.

## Results

### Sequence homology among verified effectors and the integrated prediction framework

Experimentally verified effectors were collected from literature and databases, and 653 proteins were obtained after removing redundant sequences, representing the latest and most comprehensive list of experimentally verified T4SEs[26, 43] (see Materials and Methods). Pairwise sequence alignments of full-length (FL) effector proteins or their C-terminal peptides of 100 or 50 amino acids (C100 or C50, respectively) were performed. For the FL proteins, 481 non-homologous clusters were identified after homology filtering for the proteins with > 30% identity and > 70% length coverage of the pair of proteins (FL_70%_30%_ID) (Figure 1A). However, for the C100 sequences, 249 were homologous to others with an identity of > 30%, and 473 non-redundant clusters were retained from these sequences after homology filtering (C100_30%_ID) (Figure 1A). The reduction in the number of clusters indicated that the C-terminal 100 amino acids showed more homology than the full-length effector proteins, but there were no significant differences between them (473/654 vs. 481/654, EBT *P*= 0.614). The C50 sequences further reflected the typical C-terminal homology between effectors. A total of 342 peptides were found to have homology with the others, while 401 clusters remained for these peptides after homology filtering (C50_30%_ID, 401/654 vs. 481/654, EBT *P*=3.17e-03) (Figure 1A). Rigorous homology filtering is a prerequisite for the application of machine learning to sequence analysis and effector identification. Sequence homology is often measured using similarity (SIM) rather than identity, with a cut-off of ≤ 30% for proteins. Therefore, we also employed a loose measure of homology, defined as >30% similarity, to examine sequence similarity between validated effectors. Surprisingly, the homology network involved all the 634 C100 peptides (C100_30%_SIM) (Figure 1A). The results demonstrated that the validated T4SEs showed unexpectedly significant homology, especially for the C-terminus.

**Figure 1.**
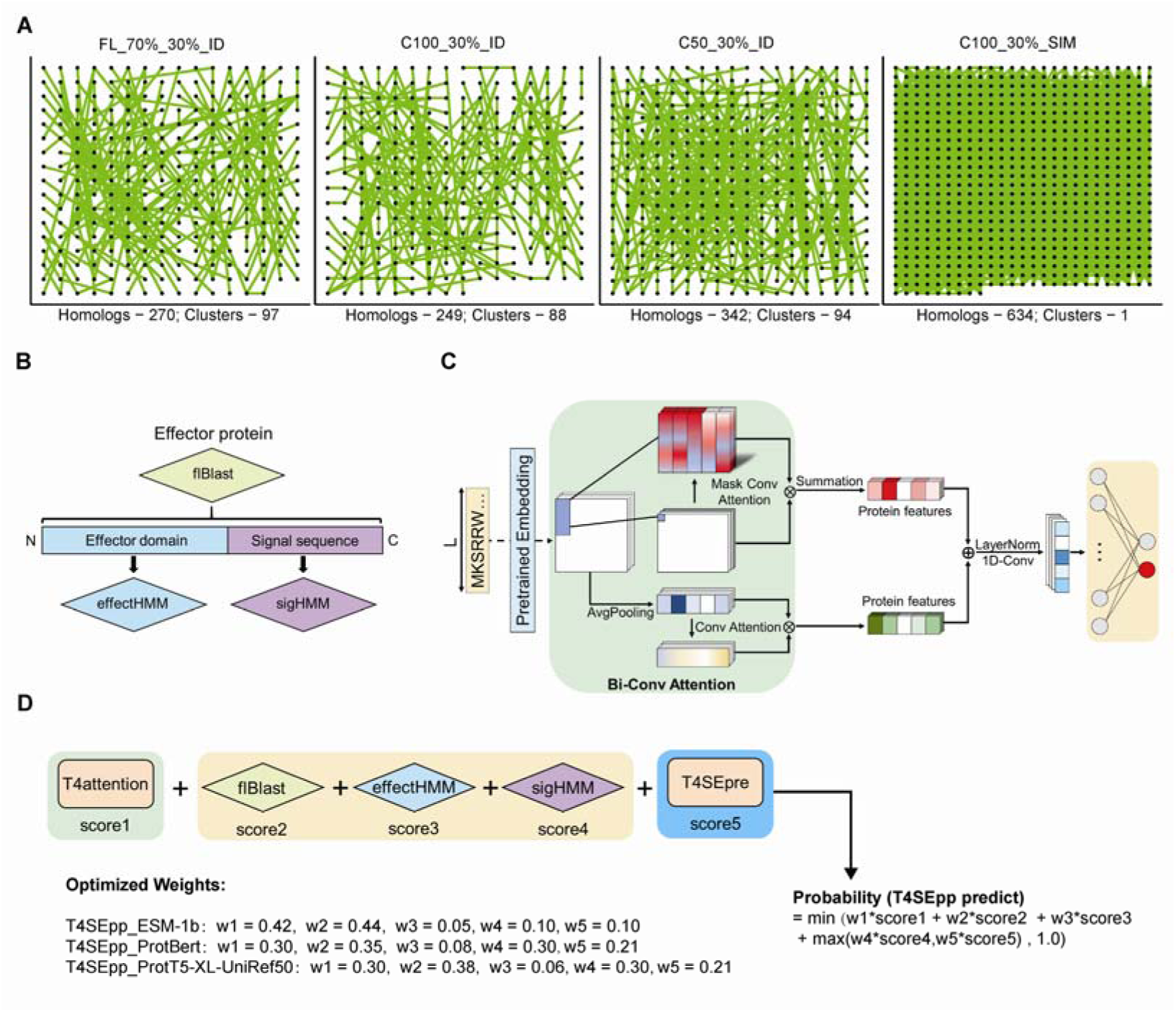
Sequence homology among T4S effectors and an integrated prediction framework. (A) Sequence homology network of T4SE. The nodes represented effectors with homology with at least one other effector. The pairs with homology (identified by the criteria defined at the top) were connected by green lines. The cluster and homology represented the number of T4SE multi-member clusters and homologous proteins. (B) Homology-based modules developed for T4SEpp, based on the full-length effector proteins (flBlast) or signal sequence (sigHMM), and effector (effectHMM) domains. (C) T4attention, a deep learning model framework based on Bi-Conv attention. (D) Flowchart of the T4SEpp prediction program. The weighted sum of the prediction scores from each individual module is incorporated into the probability that a protein is a T4SE.

Taking full advantage of the fragmental similarity between T4SEs, combined with machine learning techniques, a comprehensive prediction pipeline (T4SEpp) was designed (Figure 1B and C). Several homology searching modules have been developed to detect full-length (flBlast), effector domain (effectHMM) and C-terminal signal region (sigHMM) homologs of known T4SEs. A previous machine learning model, T4SEpre, which predicts T4SEs based on the C-terminal hand-crafted features and fine-tuned based on an updated dataset [24]. Using the generative features from pre-trained transformers, we also developed a deep learning module, T4attention, incorporated with the Bi-Conv attention mechanism. Figure 1D showes the framework of T4SEpp, taking the prediction scores of the homology search module (flBlast, effectHMM, and sigHMM), T4SEpre, and T4attention into a linear model to generate the final score, which reflects the likelihood of an input protein to be an effector.

### T4SE families of signal sequences and functional domains

According to the homology of the C50 peptides, the effectors could be clustered into 405 signal sequence families, including 94 multi-member and 311 singlet families (Supplementary Table S3). After the signal sequences (C50) were removed, 640 effectors with a length of ≥ 30 amino acids remained, of which 270 were classified into 106 multi-member families and 370 represented singlet families (Supplementary Table S4). The sequences within each multi-component family showed striking similarity, and multiple positions appeared conserved, as shown for one example, sigFAM50 (Fig. 2A). The amino acid composition (AAC) showed apparent preference in multiple positions, e.g., leucine in positions 9, 24, and 37, serine in position 18, 30, and 64, and asparagine in position 11, 26, and 48, of sigFAM50 (Fig. 2A). Effectors of the same signal sequence family may belong to different effector functional domain families and *vice versa*. For example, six cytotoxin-associated gene A (CagA) effectors and two *Legionella* proteins contained the signal sequences of the same family (sigFAM50, Figure 2B; Supplementary Table S3), but they also fell into three different effector functional domain families (effectFAM73 for all the CagAs, and effectFAM19 and effectFAM57 for the other two proteins; Figure 2B; Supplementary Table S4). This could be related to frequent domain reshuffling events that have been reported in *Legionella*[44].

**Figure 2.**
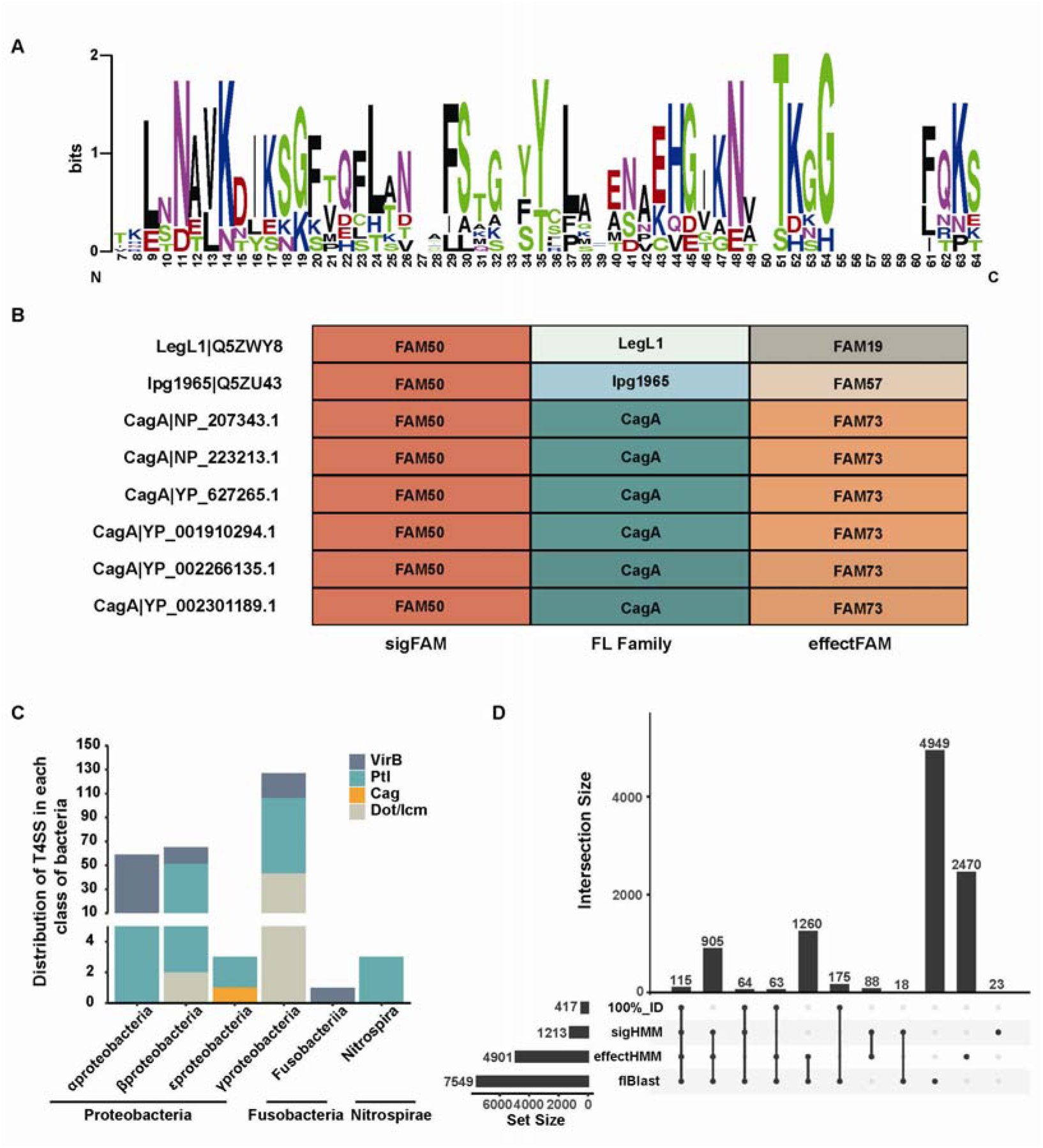
Search for T4SS and effectors in the UniProt reference proteome based on sequence homology. (A) Multiple-sequence alignment (MSA) of a homologous cluster (i.e., sigFAM50) of T4SE signal sequences. Then, utilize the sequence logo of position-specific Amino Acid Compositions (AAC) corresponding to the alignment. The height of the amino acid in each position indicated the AAC preference. (C) Using the core protein components of T4SS to construct a Hidden Markov Model (HMM) to predict the distribution of T4SS in the UniProt reference proteome. (D) Three homologous modules (sigHMM, effectHMM and flBlast) were used to predict the potential T4SE in the UniProt reference proteome containing T4SS, respectively. Where 100%_ID represents a known verified T4SE.

Furthermore, we searched for homologs of known T4SEs from the representative bacterial genomes downloaded from UniProt (8761 genomes; Supplementary Table S5). In total, 258 protein-translocating T4SSs were detected from 227 bacterial strains distributed in their phyla (*Proteobacteria*, *Fusobacteria* and *Nitrospirae*), six classes (*Alphaproteobacteria*, *Betaproteobacteria*, *Epsilonproteobacteria*, *Gammaproteobacteria Fusobacteriia*, and *Nitrospira*), 117 genera and 227 species (Figure 2C, Supplementary Table S6). In these strains with T4SSs, 10,130 proteins were detected with full-length or local homology to the known T4SEs using the individual homology searching modules, and 1,020 were identified by all the three modules (Figure 2D, Supplementary Table S7).

### Prediction of T4SEs with pre-trained transformer-based models

Recently, protein language models have been successfully applied for structural prediction and sequence classification. In this research, we used six pre-trained models, ESM-1b, ProtAlbert, ProtBert-BFD, ProtBert-UniRef100, ProtT5-XL-BFD, and ProtT5-XL-UniRef50, to generate features; based on this, we developed deep learning models (T4attention) based on Bi-Conv attention respectively to classify T4SEs and non-T4SEs. The T4attention models based on different sequence embedding features were compared for performance based on a five-fold cross-validation strategy (Table 1). Generally, T4attention_ESM-1b performed the best, followed by T4attention_ProtT5-XL-UniRef50, and T4attention_ProtAlbert showed the poorest performance, according to the Matthew’s correlation coefficient (MCC) and F1-score (Table 1). T4attention_ESM-1b not only reached the highest MCC and F1-score (0.861 and 0.819, respectively), but required the lowest computational resources (Supplementary Figure S3). It was also noted that, for the same protein language model architecture, ProtBert or ProtT5-XL, for example, the generation of features from models pre-trained from various volumes of protein database required similar computational resource, but the smaller database-based pre-trained models always generated features for subsequent T4attention models with better performance (MCC of T4attention_ProtBert vs. T4attention_ProtBert-BFD, 0.814 vs. 0.797; T4attetion_ProtT5-XL-UniRef50 vs. ProtT5-XL-BFD, 0.818 vs. 0.800) (Table 1, Supplementary Figure S3). The redundancy of protein sequences in the BFD dataset might lead to biases in model training, and further compromise the performance of models addressing downstream tasks.

**Table 1.**
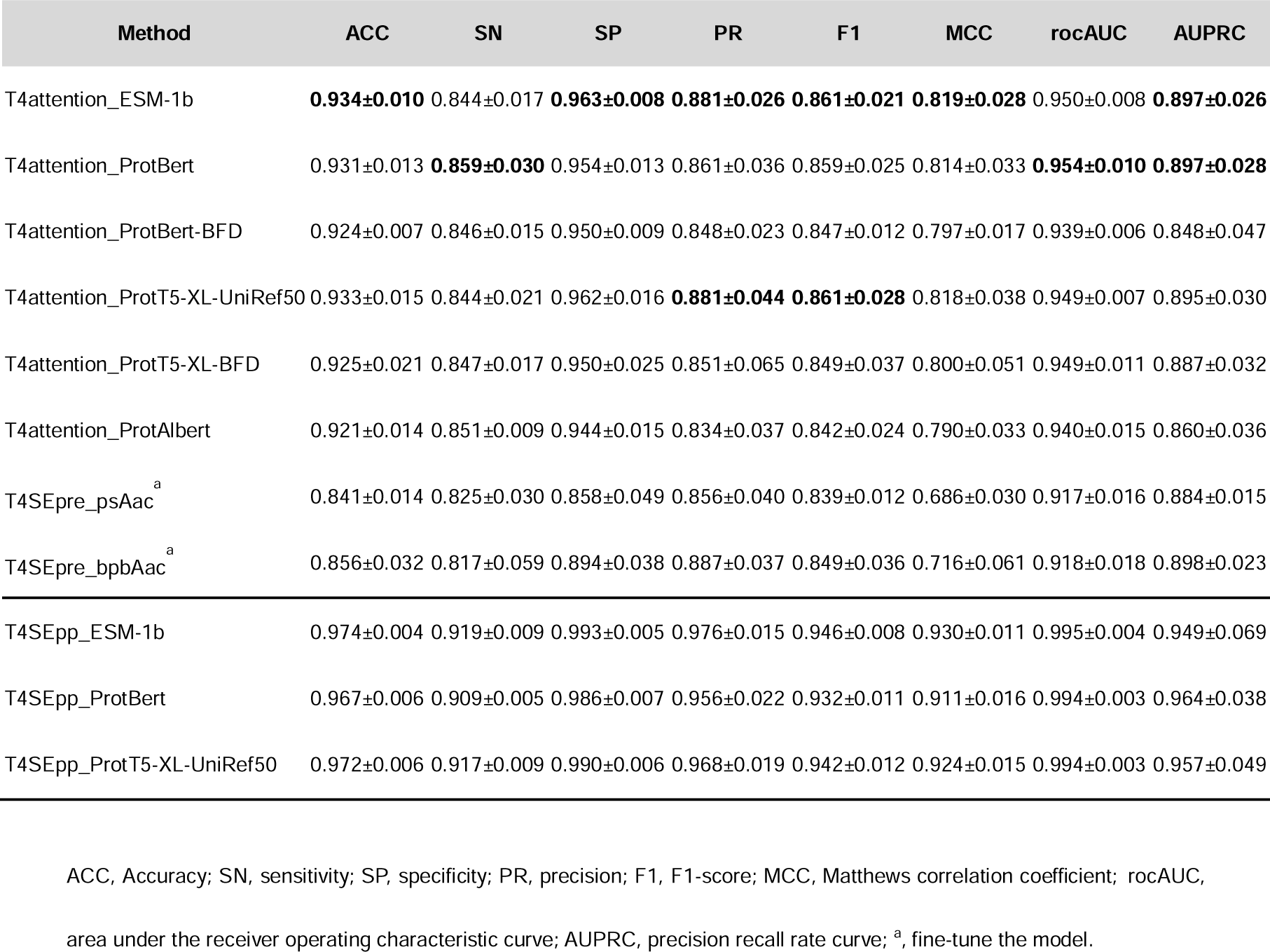
Performance comparison of the models in T4SEpp on 5-fold cross-validation dataset.

We also evaluated the performance and generalization abilities of these models on an independent testing dataset. T4attention_ProtBert showed the overall the best performance, for which the MCC, F1-score, and accuracy reached 0.917, 0.927, and 0.987, respectively (Table 2). T4attention_ESM-1b was unexpected and showed poor performance (Table 2). Consistent with the cross-validation results, the ProtBert and ProtT5-XL models, based on the features generated by transformers pre-trained from a smaller database (UniRef100/UniRef50), showed better performance (Table 2, Supplementary Figure S4).

**Table 2.**
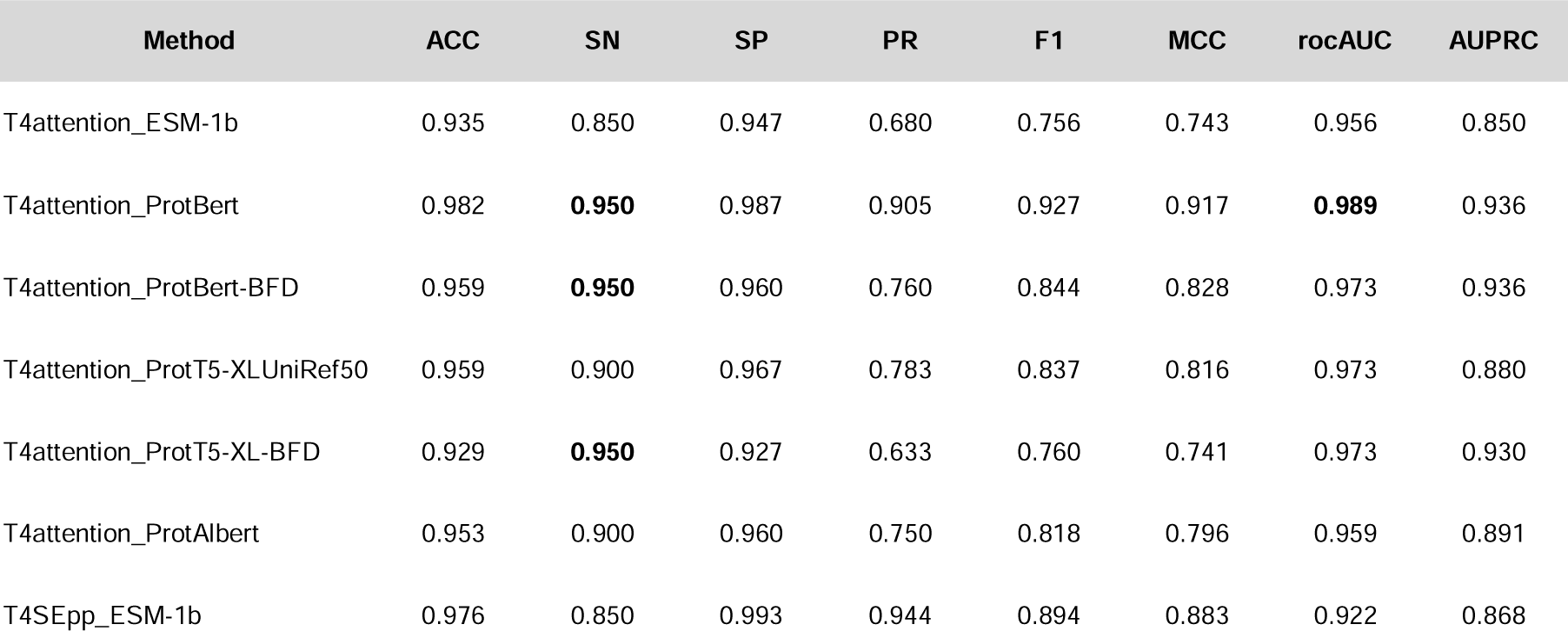
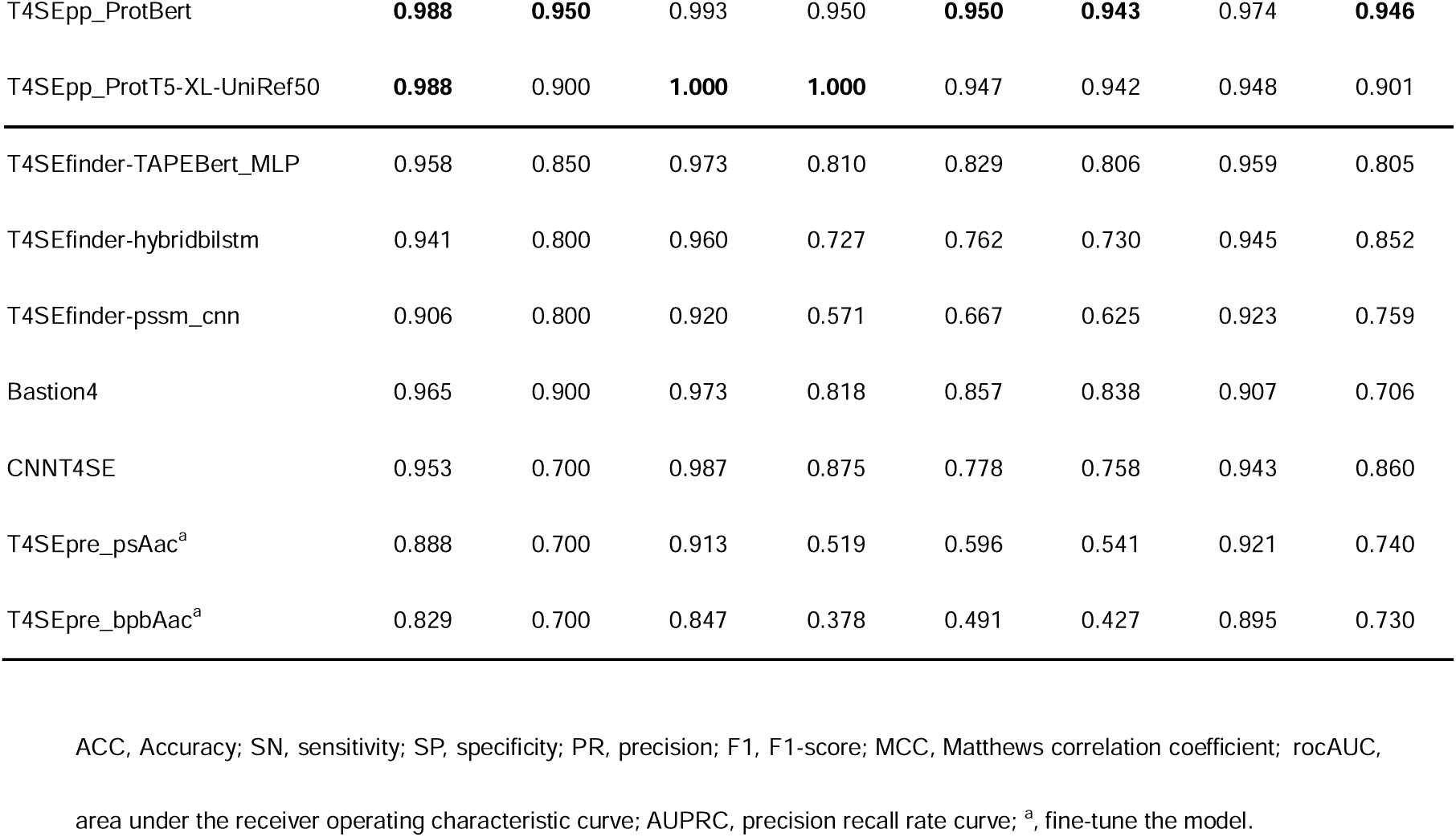
Performance comparison of the models in T4SEpp and other tools on the independent dataset.

Considering the performance of models based on both cross-validation results and the independent testing dataset, as well as the requirement of computational resources, we integrated three models, T4attention_ESM-1b, T4attention_ProtBert, and T4attention_ProtT5-XL-UniRef50, into the pipeline to predict T4SEs.

### An integrated pipeline predicting T4SEs with largely improved performance

In addition to the models based on the features generated by the transformer, we tested traditional machine learning models based on hand-crafted features. To this end, we fine-tuned two models of T4SEpre models (T4SEpre_psAac and T4SEpre_bpbAac) to learn the amino acid composition features in the C-termini of T4SEs[24]. Both models showed a certain performance in the prediction of T4SEs according to the cross-validation results or the independent testing dataset, although they were not comparable to the T4attention models (Tables 1 and 2).

To further improve the accuracy and reduce the false positive rate for T4SE prediction, we assembled a unified pipeline, T4SEpp, integrating the homology searching modules, machine learning models based on hand-crafted features and models based on transformer-generated features (Figure 1). The integrated pipeline showed strikingly better performance than the individual models, with MCC values of 0.930, 0.911 and 0.924 for T4SEpp_ESM-1b, T4SEpp_ProtBert, and T4SEpp_ProtT5-XL-UniRef50 based on the cross-validation evaluation and 0.883, 0.943, and 0.942 for the testing dataset, respectively (Tables 1 and 2).

T4SEpp was also compared to other state-of-the-art(SOTA) T4SE prediction models, such as Bastion4[26], CNNT4SE[27] and T4SEfinder[29]. Among these other models, Bastion4 showed the best performance, which was close to that of the T4attention models but was far inferior to the integrated T4SEpp (Table 2).

### Genome-wide screening of T4SEs in *Helicobacter pylori* and other bacteria

*H. pylori* is a gram-negative, spiral-shaped bacterium that colonizes the stomach in approximately half of the world’s population[45]. Although most individuals do not experience any adverse health outcomes attributable to *H. pylori*, the presence of these bacteria in the stomach increases the risk of developing gastric diseases[46-50]. *H. pylori* infection is also the strongest known risk factor for gastric cancer, the third leading cause of cancer-related death worldwide[51]. T4SS plays an important role in *H. pylori*[47-50]. However, to date, only one T4SE, CagA, has been identified for the T4SS in *H. pylori*[52]. Here, we applied T4SEpp to screen T4SE candidates from the proteins derived from the genome of *H. pylori* 26695, a model *H. pylori* strain (NCBI accession number: NC_000915.1). The three T4SEpp integrated models, T4SEpp_ESM-1b, T4SEpp_ProtBert, and T4SEpp_ProtT5-XL-UniRef50, predicted 55, 22, and 38 T4SE candidates, respectively, and 13 were shared by the prediction results of all the three models (Figure 3A-B; Supplementary Tables S6, S8). The 13 potential effector genes were scattered throughout the genome (Figure 3B). Notably, *HP_RS02695,* which encodes the only known effector CagA, was among the 13 candidates (Figure 3B).

**Figure 3.**
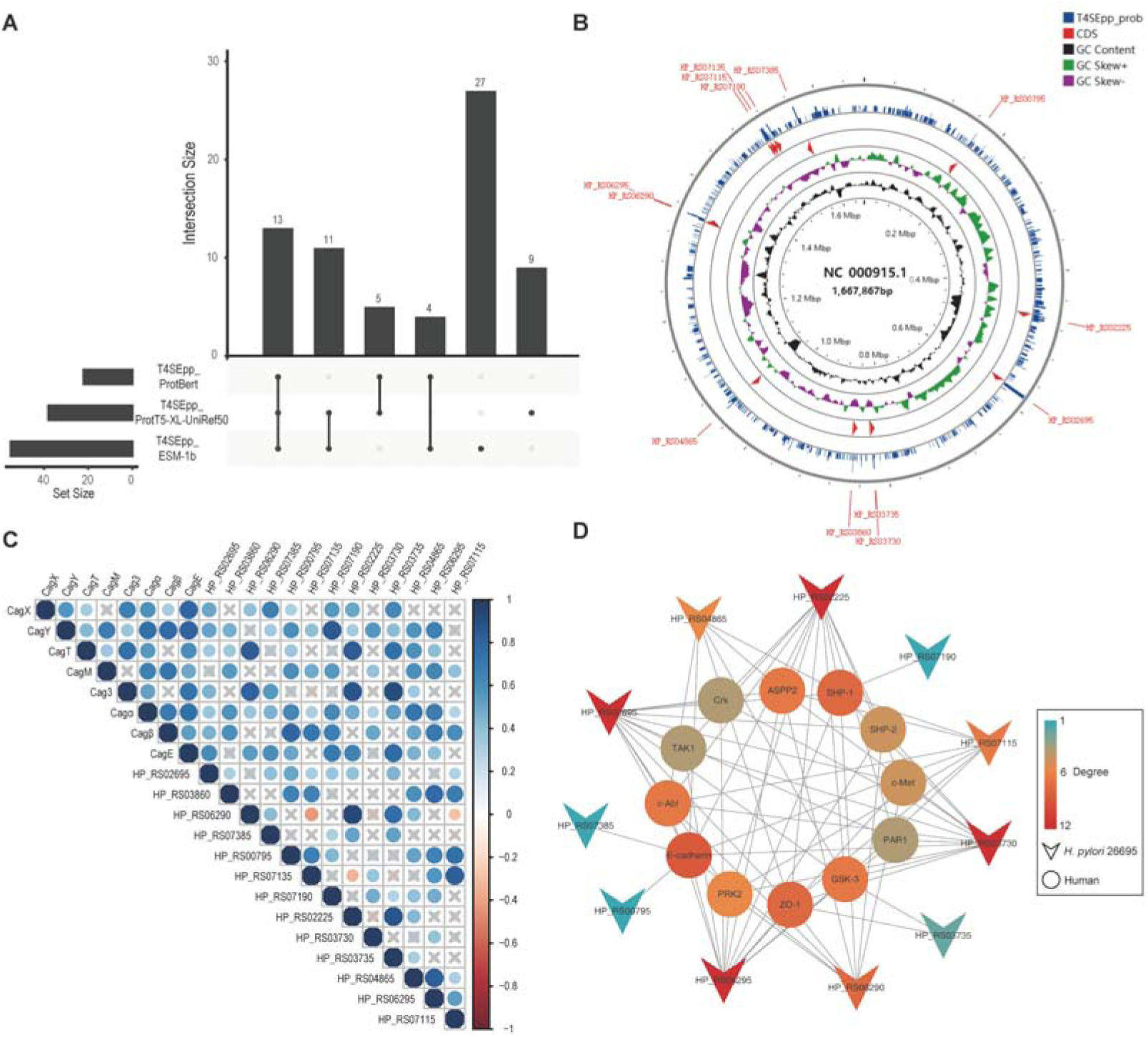
Whole-proteome detection for T4SEs in pathogenic bacteria (*H. pylori* 26695). (A) Prediction of potential T4SEs in the *H. pylori 26695* proteome using three T4SEpp models. (B) Use the circos diagram to show the distribution of potential T4SEs predicted by the three T4SEpp models on the *H. pylori 26695* chromosome (NC_000915.1), where T4SEpp_prob represents the mean value of the prediction results of the three T4SEpp models, and the outer circle of the circos diagram represents the three T4SEpp model predictions were all positive. (C) Under 12 different expression conditions of *H. pylori 26695*, the expression correlation of Cag T4SS core components with 12 potential T4SEs and CagA (HP_RS02695) predicted by three T4SEpp models were positive. (D) Prediction of potential interactions between 12 potential T4SEs in H. pylori 26695 and 12 human proteins using DeepHPI. These 12 human proteins are known to interact with CagA(HP_RS02695).

Gene co-expression was analyzed for the 13 T4SE candidate genes in *H. pylori 26695* using an RNA-seq dataset sampled from the strain collected under 12 different conditions[53]. Except for *HP_RS06290*, *HP_RS03730*, *HP_RS04865*, and *HP_RS06295*, the remaining eight genes showed a strong expression correlation with *cagA* expression (Figure 3C). The genes co-expressed with *cagA* also showed a significant correlation with the expression of the core component genes of the Cag T4SS (Figure 3C). Furthermore, we annotated 12 human proteins that showed experimentally verified interactions with CagA by literature search, including ASPP2, c-Abl, c-Met, Crk, E-cadherin, GSK-3, PAR1, PRK2, SHP-1, SHP-2, TAK1, and ZO-1[54-65]. The interaction network between the 13 potential *H. pylori* 26695 T4SEs and the 12 human proteins was inferred (Figure 3D). Ten of the candidate T4SEs showed potential interaction with at least one of the human proteins (Figure 3D). Similar to CagA, HP_RS02225, HP_RS06295 and HP_RS03730 showed interacted with all the 12 human proteins (Figure 3D). Taken together, the proteins predicted by T4SEpp could potentially represented new T4SEs, or may be closely related to the pathogenicity of *H. pylori 26695*.

We also used T4SEpp to screen the T4SE candidates from the genomes of 227 bacterial strains bearing T4SSs. T4SEpp_ESM-1b, T4SEpp_ProtBert, and T4SEpp_ProtT5-XL-UniRef50 detected 16,972, 20,441 and 17,197 T4SE candidates respectively, with 12,622 common candidates co-predicted by all the three T4Sepp models (Supplementary Table S9, Supplementary Figure S5).

### Web server and implementation of T4SEpp

To facilitate the implementation of T4SEpp, we developed a user-friendly web application (https://bis.zju.edu.cn/T4SEpp). The three T4SEpp integrated models, T4SEpp_ESM-1b, T4SEpp_ProtBert, and T4SEpp_ProtT5-XL-UniRef50 can be chosen and implemented by users. Both the overall prediction results and the results of the individual modules are displayed in table format, which can be downloaded and filtered easily.

## Discussion

T4SS plays a crucial role in bacterial pathogenicity by secreting effectors into host cells. *L. pneumophila* can translocated more than 300 known effectors into human cells via the Dot/Icm T4SS system, causing legionellosis[66, 67]. In *H. pylori*, CagA is the only known T4SE that can hijack multiple signaling pathways in gastric epithelial cells, leading to gastritis, gastric ulcer and even gastric cancer[68, 69]. Identifying the full repertoire of T4SEs in a pathogen is important to understand its pathogenic mechanisms. Computational methods can assist with the effective identification of new effectors[70]. However, the currently available T4SE prediction tools still show high false positive rates[2]. To address this issue, we developed a unified T4SE prediction pipeline, T4SEpp, which includes homologous search modules, traditional machine learning modules and natural language processing-based modules. T4SEpp outperformed other SOTA methods for predicting T4SEs, with improved sensitivity and specificity. Furthermore, we initiated a web server that can conveniently implement the T4SEpp pipeline, providing the prediction results for each module.

Although the component modules of T4SEpp can be used for T4SE prediction, they often show higher false positive rates when used alone. This could be related to the low power of the individual dimensions of the features. Specifically, T4SE signal sequences were considered to contain important common features guiding T4SE secretion and translocation, which were used for effective T4SE prediction using tools such as T4SEpre[24]. However, the computational models based only on the signal sequences showed performance inferior to other models based on multiple-aspect features extracted from full-length proteins[26]. In this study, we discovered high sequence similarity in the C-terminal signal region among the proteins, without apparent homology to full-length effectors. Such undetected homology could have introduced bias and led to overfitting of various established machine learning algorithms and the discrepancy between the reported and actual accuracy of these methods. However, the C-terminal homology could also suggest the independent evolution of the signal sequences, and it could potentially be applied to facilitate the identification of new effectors[42].

In this study, three types of modules were integrated to predict T4SEs. Homology searching-based modules provide more accurate results, but they also show a lower capacity to detect new effectors with or without remote homology. The re-trained T4SEpre modules focused on the important features of the C-terminal signal sequences of T4SEs. T4attention learns from the full-length effector proteins the features generated by protein language models (pLMs) pre-trained with large-scale protein databases. These pLM-based models can learn new, previous unknown features that may involve position-position interactions, and have demonstrated outstanding performance in the prediction of proteins with various biological functions, such as subcellular localization and secondary structure. We used multiple pLMs to build transfer learning models, most of which exhibited excellent performance in T4SE prediction. Interestingly, we noticed that the pre-trained pLMs based on the larger datasets did not generate better prediction performance. pLMs pre-trained on smaller datasets are more efficient. Therefore, the transfer models were trained with the pLMs based on smaller non-redundant protein datasets. T4SEpp, which integrated all three types of modules, significantly outperformed both individual modules and other similar applications.

Using T4SEpp, we analyzed the potential new T4SEs in both *H. pylori* and other strains bearing T4SS. We identified 12 new T4SEs in *H. pylori*. We also identified 12,205 new T4SEs and 417 known T4SEs from 227 strains bearing a T4SS. The results suggested that there are many new effectors yet to be clarified.

Despite the significant performance improvement of T4SEpp, there remains a need to further improve the prediction of T4SEs. Other features that have been known to contribute to the recognition of T4SEs, such as the GC content of genomic loci, phylogenetic profiles, consensus regulatory motifs in promoters, physicochemical properties, secondary structures, homology to eukaryotic domains, and organelle-targeting signals, have not been integrated into the current version of the model[70]. Novel features that could be further integrated to improve the model performance remain to be disclosed. The different types (IVA and IVB) of effectors, chaperone-dependent or chaperone-independent effectors, or species-specific effectors can also be modeled and predicted separately to make more accurate prediction[70].

## Materials and methods

### Datasets

The 390 T4SEs used by Bastion4 as the positive training dataset[26] and 540 T4SEs annotated in SecReT4 v2.0[43] were collected and merged, and in total we got 653 non-identical, validated T4SEs. CD-HIT[71] was used to filter homology-redundant proteins with sequence identity ≥ 60%, generating 518 non-redundant T4SEs, which were used as the positive training dataset(Supplementary Figure S1A). For the negative training dataset, we collected 1112 and 1548 non-T4SE protein sequences from Bastion4[26] and PredT4SE-stack[72], respectively. The same procedure was used to eliminate the sequence redundancy among the non-T4SEs and between the non-T4SEs and T4SEs in the positive training dataset, generating 1590 non-redundant non-T4SEs (Supplementary Figure S1A). An independent validation dataset was also prepared, for which the T4SEs were collected from the testing dataset of Bastion4 (30) and others (74) annotated from literature published recently (Supplementary Table S1), and the 150 testing non-T4SEs of Bastion4 were also used as negative ones. CD-HIT was used to filter the redundant proteins with ≥60% sequence identity to the training proteins and among proteins in the validation dataset, resulting in 20 non-redundant T4SEs and 150 non-T4SEs (Supplementary Figure S1B).

### Genome-wide screening of protein-translocation T4SSs

The conserved core component proteins were collected from four representative protein-translocation T4SSs, including the *Agrobacterium tumefaciens* VirB/VirD4 T4SS (inner membrane complex proteins VirB3, VirB6, VirB8, VirB10 and VirD4, and outer membrane complex proteins VirB7, VirB9 and VirB10)[16], the *Bordetella pertussis* Ptl T4SS (inner membrane complex proteins PtlB, PtlE and PtlH, and outer membrane complex proteins PtlF and PtlG)[73], the *Helicobacter pylori* Cag T4SS (inner membrane complex proteins Cagα, Cagβ and CagE, and outer membrane complex proteins CagX, CagY, CagT, CagM and Cag3)[18], the *Legionella pneumophila* Dot/Icm T4SS (inner membrane complex proteins IcmB, IcmG and DotB, and outer membrane complex proteins DotC, DotD, DotG and IcmK)[16]. Hidden Markov Model (HMM) profiles were built using HMMER 3.1 for the T4SS component protein families[74]. Protein sequences derived from the 8761 reference bacterial genomes curated in UniProt were scanned with HMMER and the HMM profiles to determine the distribution of homologs of T4SS core component proteins (Supplementary Table S5).

### Homology networks of the T4SE peptide sequences

The sequences of 653 non-identical verified T4SE proteins were used to construct the homology networks. JAligner implemented the Smith-Waterman algorithm to determine the similarity between any pair of full-length effectors or peptide fragments of designated length (http://jaligner.sourceforge.net/). The identity and similarity percentages between any pair of sequences were used as measures to determine the homology level[38].

### Homology-based T4SE detection modules

Diamond blastp was used to determine the homology and cluster the full-length effector proteins[75] and to screen new full-length homologs (flBlast). Two proteins showing ≥30% similarity for ≥70% of the full length of either protein were considered to be full-length homologs[38, 76]. The C-terminal 50-aa signal sequences of the verified effectors were clustered according to homology networks with 30% identity for 70% length aligned by JAligner. HMM profiles were built for each signal sequence family, and a sigHMM module was developed to screen for proteins with C-terminal sequences homologous to the profiles of known T4SE signal sequence families. The homology cutoff for HMM searching was optimized for each family, ensuring that all or most of the known effectors recalled and maintained a higher specificity. For effectHMM, we removed the C-terminal 50-aa signal from each known effector sequence, and the remaining peptide fragment with >30-aa length was used for domain clustering. Pairwise alignment was repeatedly performed with BLAST between the domain sequences, and the cutoff for homology was optimized based on the average coverage of the aligned length multiplied by the identity, that is, ≥10[38]. The HMM profiles were built for the effector domain families, and effectHMM was developed using a similar procedure as sigHMM to screen the proteins with homologous T4SE effector-domains. We used EBT to compare general homology between proteins[38, 77].

### Fine-tune T4SEpre models with updated datasets

Fine-tune T4SEpre models (T4SEpre_psAac and T4SEpre_bpbAac) using the new training datasets of T4SEs and non-T4SEs. The original T4SEpre procedure was followed for feature representation, parameter optimization and model training[24]. Briefly, sequential amino acid, bi-residue and motif composition features and position-specific amino acid composition profile for the positive training dataset were represented for each C-terminal 100-aa sequence for the psAac model. For the bpbAac model, position-specific amino acid composition profiles of both the positive and the negative training datasets (Bi-Profile Bayesian features) were represented for each C-terminial 100-aa sequence. Support vector machine (SVM) models were trained for feature matrices. The kernel functions, that is, linear, polynomial, sigmoid, and radial base function (RBF), and corresponding parameters (cost and gamma) were optimized using a 5-fold cross-validation grid search strategy. The sklearn v1.0.1 was used for implementing SVM model training and kernel/parameter optimization.

### The deep learning architecture of T4attention based on pre-trained protein language models

#### Input embeddings

Frozen embeddings were extracted directly from protein language models (pLMs) without fine-tuning the training data. Four different basic LMs were used in this study, and six different pLMs were pre-trained with different datasets. The basid LMs include, (i) “ESM-1b”[33], which is a Transformer model, (ii) "ProtBert" [32], which is a BERT-based encoder model[30], generating two pLMs pre-trained on BFD[78] and UniRef100[79] data, respectively, (iii) ProtT5-XL[32], which is an encoder model based on T5[80], generating two pLMs pre-trained on BFD and UniRef50, respectively, and (iv) ProtAlbert[32], which is an encoder model based on Albert[81] and pre-trained only with UniRef100.

#### Optimization strategy

We use a BERT-like optimizer AdamW and a Cosine Warm-up strategy[30] to optimize the loss of the learning model. The initial learning rate is set to 0.0001, the batch size is set to 18, and the warm-up steps were set to 10. An early stopping strategy was applied to monitor the validation ACC with 30 epochs to prevent overfitting. To address the challenges of imbalanced positive and negative samples and the difficulty of training individual samples in deep learning model training, we adopted the Focal Loss method to mitigate the issue of gradient descent difficulty[82]. Focal Loss increases the hyperparameter γ (default γ=2) based on the weighted cross-entropy loss, which controls the shape of the curve.

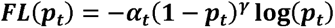

*α_t_*: Weight of the sample t,

*p_t_*: Binary cross entropy loss.

#### T4attention model

The input to T4attention (Figure 1C, Supplement Figure S2) is a protein embedding ***E***_0_ ϵ ℝ^n×d_0_^, where **n** is the sequence length and **d_0_** is the size of the embedding (depending on the feature extraction model). T4attention is a model based on Bi-Conv attention. In the protein embedding direction, average pooling is performed directly, and the input is transformed by two separate 1D convolutions, where the 1D convolution serves as the attention coefficient **e** and value **v** for computing the embedding dimension, *e, v* ϵ ℝ^d_1_^. Thus, we obtained the feature representation of the embedding dimension ***x = softmax(e) x v***. In the direction of the protein sequence, we randomly intercept the length of **m** in the length direction of the protein-embedding sequence such that the protein embedding becomes *E*_1_ ϵ ℝ^m×d_0_^. Similar to the convolutional attention calculation in the protein embedding direction, the attention coefficient **e’** and value **v’** are obtained, e’, v’ ϵ ℝ^m×d_1_^. The difference is that the direction of the convolution is in the direction of the sequence length, so that we can obtain the feature representation of the protein sequence direction and converge according to the sequence length direction by 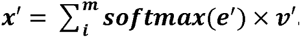. The convolution attention results of the embedding direction and the protein sequence direction are merged and passed through the LayerNorm and the residual one-dimensional convolution, and the class probabilities are obtained through the two-layer multi-layer perceptron (MLP), ***p*(c|x) = *softmax(MLP*(*Conv*(*x* + *x*^′^) + (*x* + *x*^′^)))**, where **c** indicates the category of the output (i.e., T4SE or nonT4SE).

T4attention was developed using PyTorch v1.10.1. The models were trained and evaluated with 24-GB of memory and an NVIDIA GeForce RTX 3090 GPU for acceleration.

### Integrated T4SE prediction model

T4SEpp is a linear model that integrates multiple prediction modules developed or re-trained in this study, including homology-searching modules for full-length or fragmented effector proteins, traditional machine-learning modules with hand-crafted features, and the attention-based transfer learning modules using the features generated by pre-trained protein language models. For any prediction module, the factor was set to 1.0 if there was a positive prediction result, and 0 otherwise. Weight **x** was assigned empirically to each module, where x ϵ (0,0.50). The maximum T4SEpp predicted value was set as 1.0. We trained the model using a grid search with 5-fold cross-validation to determine the optimal combination of weights. The early stopping strategy was similar to that used for T4attention. The final optimal parameters were shown in Figure 1D.

### Assessment of model performance

Measures including accuracy (ACC), sensitivity (SN), specificity (SP), precision (PR), F1-score, Matthew’s correlation coefficient (MCC), the area under the receiver operating characteristic curve (rocAUC), and the precision recall rate curve (AUPRC) were calculated to evaluate and compare the performance of models predicting T4SEs. Some of these measures are defined as follows:

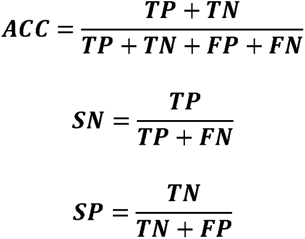

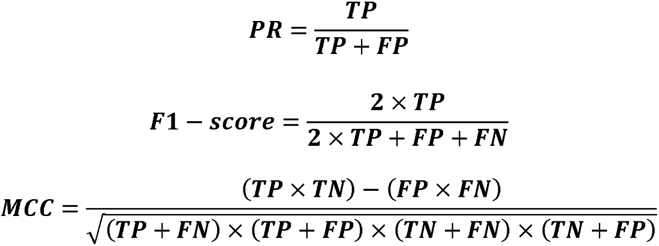

where TP, TN, FP, and FN represent the number of true positives, true negatives, false positives, and false negatives, respectively.

### RNA-seq analysis

RNA-seq datasets of *H. pylori* 26695 under different conditions were downloaded from the NCBI GEO DataSets database with accessions GSE165055 and GSE165056[53]. After removing the adapters and low-quality sequences with Trimmomatic v0.39[83], the cleaned reads were mapped to the *H. pylori* 26695 reference genome (NC_000915.1) using READemption (Version 2.0.0)[84]. The annotated genes were then quantified and analyzed. Protein-Protein Interaction (PPI) Networks were built and visualized using the Cytoscope v3.9.1[85].

## Availability

The online version of the T4SEpp is freely accessible at https://bis.zju.edu.cn/T4SEpp. The standalone version of the T4SEpp model and the individual modules were are also deposited at https://github.com/yuemhu/T4SEpp. RNA-seq data are publicly available in the NCBI GEO DataSets database with accession numbers GSE165055 and GSE165056.

## Authors’ Contribution

MC conceived and supervised the project. YH, MC, and YW coordinated the project. YH, YZ, YH, and ZZ dataset collection. YH provided codes, models and software tools. YH, XH, and HC developed the website. YH and YW performed model comparison and RNA-seq data analyses. YH, XH, HC, SL, QN, YW, and MC wrote the first draft of this manuscript. YH, YW, and MC revised the manuscript accordingly.

## Supplementary data

**Supplementary Figure S1.** The workflow to construct the training(A) or independent testing(B) dataset in this study.

**Supplementary Figure S2.** Two modules used by the T4attention model.

**Supplementary Figure S3.** The relationship between the feature extraction time of 6 different protein natural language models and the prediction performance of T4attention model F1-score (A) and MCC (B) in the 5-fold cross-validation dataset.

**Supplementary Figure S4.** The relationship between T4attention model prediction performance F1-score (A) and MCC (B) in the independent test set and the overall time-consuming use of 6 different protein natural language models to extract features and their T4attention model predictions.

**Supplementary Figure S5.** Three T4SEpp model were used to predict the potential T4SE in the UniProt reference proteome containing T4SS, respectively. Where 100%_ID represents a known verified T4SE.

**Supplementary Table S1.** The 74 T4SEs independently collected from the literature.

**Supplementary Table S2.** Hyperparameters used in deep learning models of T4attention.

**Supplementary Table S3.** Homologous Clusters of T4S Effector Signal Sequences.

**Supplementary Table S4.** The distribution of effector domain families.

**Supplementary Table S5.** Distribution of the Uniprot Bacteria Reference Proteomes (Download date October 19, 2022).

**Supplementary Table S6.** Distribution of T4SS in the UniPort bacterial reference proteome.

**Supplementary Table S7.** Homology prediction results of T4SE in strains containing T4SS in the Uniport Bacteria Reference Proteomes.

**Supplementary Table S8.** Distribution of potential T4SEs in the *H. pylori_26695* (NC_000915.1).

**Supplementary Table S9.** Distribution of potential T4SEs in the Uniport Bacteria Reference Proteomes.

