## Supplementary figures and images for "T4SEpp: a pipeline integrated with protein language models effectively predicting bacterial type IV secreted effectors" (from another version of this preprint: version 1, 2023-07-03)

### Supplementary Figure S1

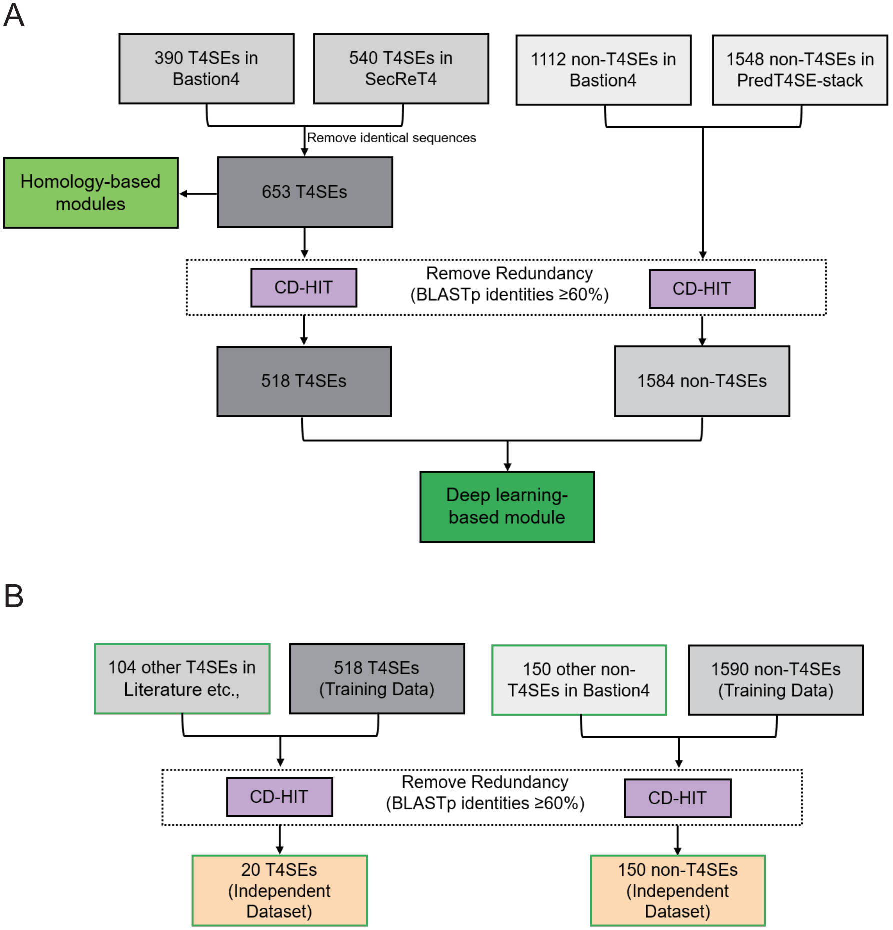

### Supplementary Figure S2

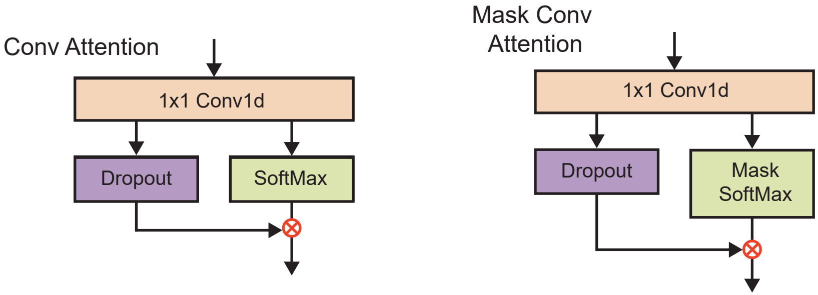

### Supplementary Figure S3

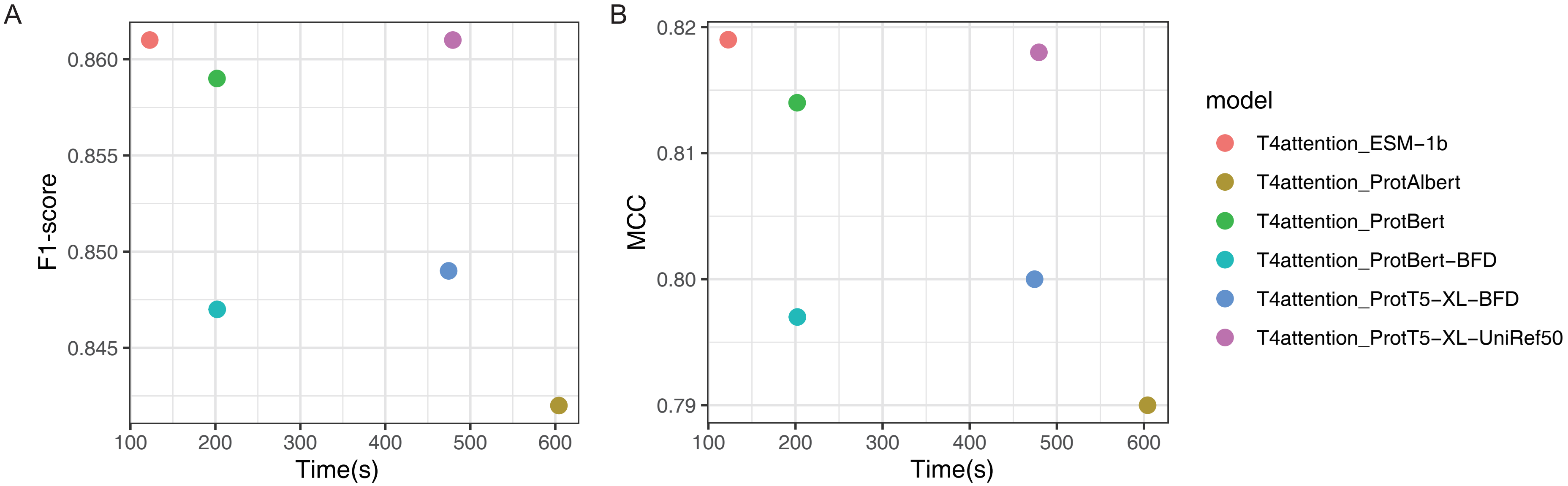

### Supplementary Figure S4

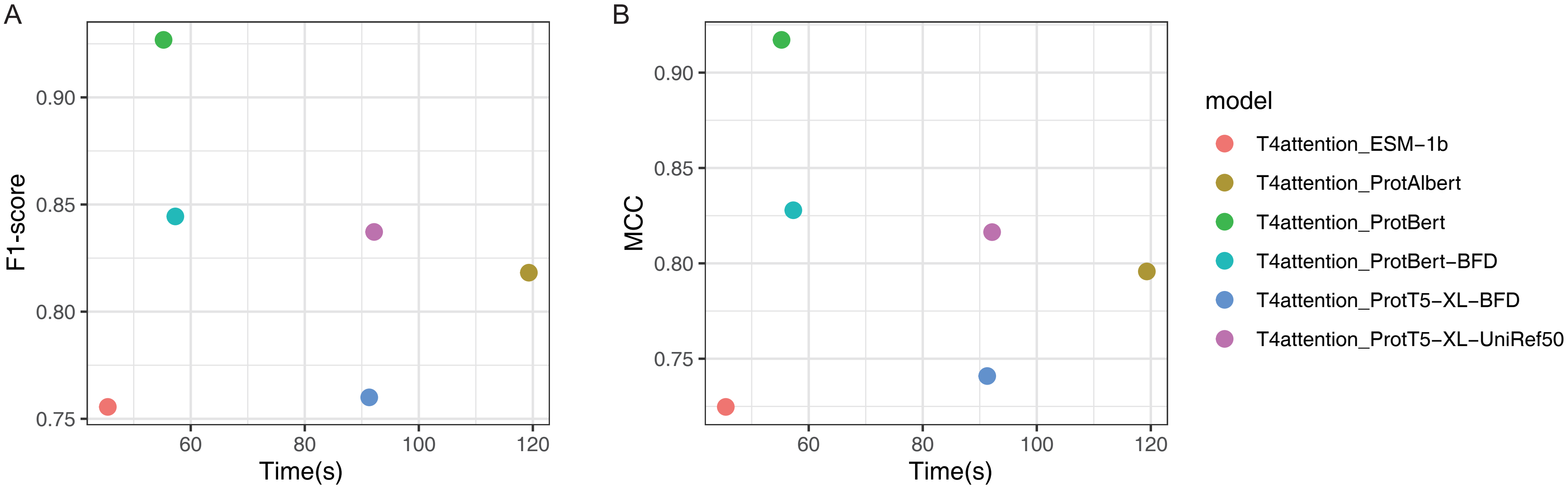

### Supplementary Figure S5

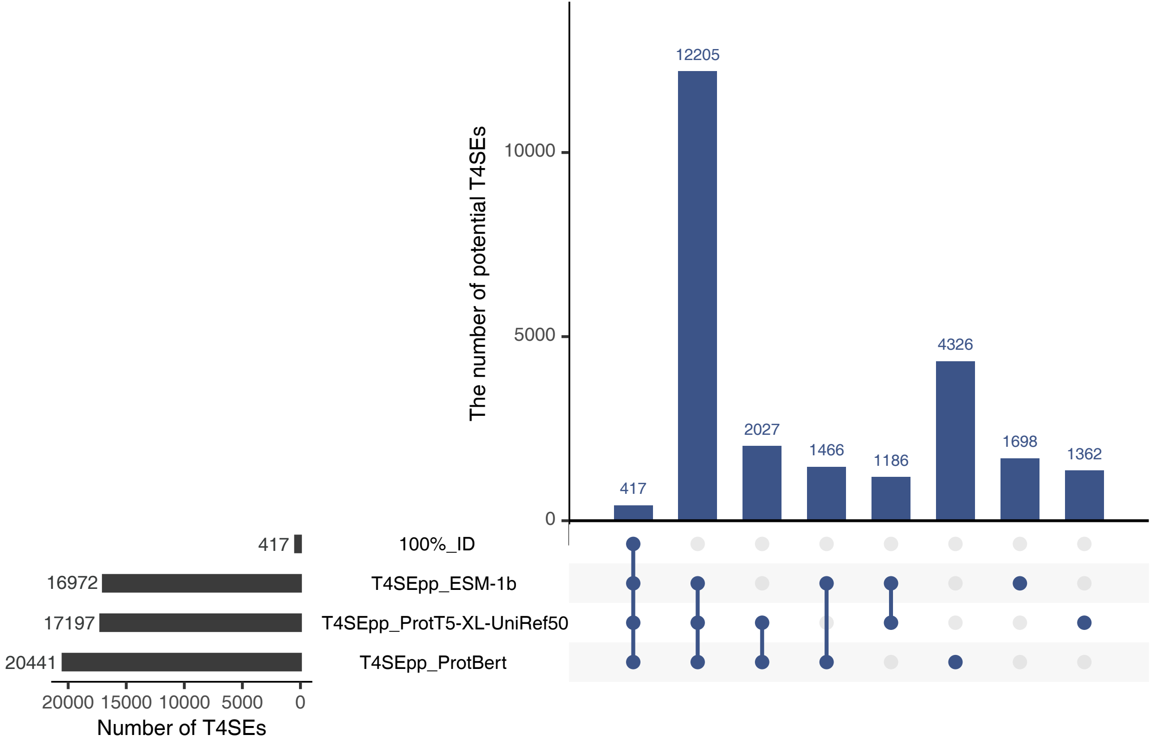
